# Rapid genome recoding by iterative recombineering of synthetic DNA

**DOI:** 10.1101/115493

**Authors:** Yu Heng Lau, Finn Stirling, James Kuo, Michiel A.P. Karrenbelt, Yujia A. Chan, Adam Riesselman, Connor A. Horton, Elena Schaefer, David Lips, Matthew T. Weinstock, Daniel G. Gibson, Jeffrey C. Way, Pamela A. Silver

**Author notes:** **Corresponding author:** Pamela A. Silver, Department of Systems Biology, Harvard Medical School, 200 Longwood Avenue, Alpert 536, Boston, MA 02115, USA.

## Abstract

Genome recoding will provide a deeper understanding of genetics and transform biotechnology. We bypass the reliance of previous genome recoding methods on site-specific enzymes and demonstrate a rapid recombineering based strategy for writing genomes by Stepwise Integration of Rolling Circle Amplified Segments (SIRCAS). We installed the largest number of codon substitutions in a single organism yet published, creating a strain of *Salmonella typhimurium* with 1557 leucine codon changes across 200 kb of the genome.

---

The next widely anticipated breakthrough in genetic engineering is the ability to rapidly rewrite the genomes of industrially relevant microbes, plants, and animals. Rewriting entire genomes will deepen our understanding of the genetic code and dramatically transform human health, food and energy production, and our environment^1–5^. A major challenge identified by the Genome Project-Write consortium is the efficiency of building and testing large modified genomes^1^. In particular, genome recoding involves synonymous replacement of all instances of specific codons throughout an entire genome^2^, requiring efficient assembly of large constructs containing thousands of designed base changes^3^. New foundational technologies are therefore crucial for accelerating the pace of genome synthesis and modification.

No effective strategy has previously been demonstrated for recoding large contiguous genomic regions with high frequency codon substitutions. Recent efforts in *Escherichia coli* recoded independent 20-50 kb genomic regions, using site-specific integrases^3^ or Cas9 endonucleases^6^ that require additional time-consuming cloning steps. Alternative editing methods such as multiplex automated genome editing^7,8^ are also not sufficiently high throughput when thousands of codon changes must be made. Landmark work on constructing a synthetic yeast genome^4^ is enabled by the high efficiency of native homologous recombination which is specific to yeast, and does not apply to most other organisms.

Our rapid genome recoding approach leverages direct iterative recombineering^9,10^ and bypasses the reliance on site specific enzymes. In this work, we use Stepwise Integration of Rolling Circle Amplified Segments (SIRCAS) to accumulate 1557 codon changes in 176 genes across 200 kb of the *Salmonella enterica*serovarTyphimurium LT2 genome (Figure 1). This strain contains the largest number of codon substitutions in a single recoded organism published to date. A recoded designer *S. typhimurium* could have diagnostic and therapeutic applications in the human body^11–13^.

**Figure 1.**
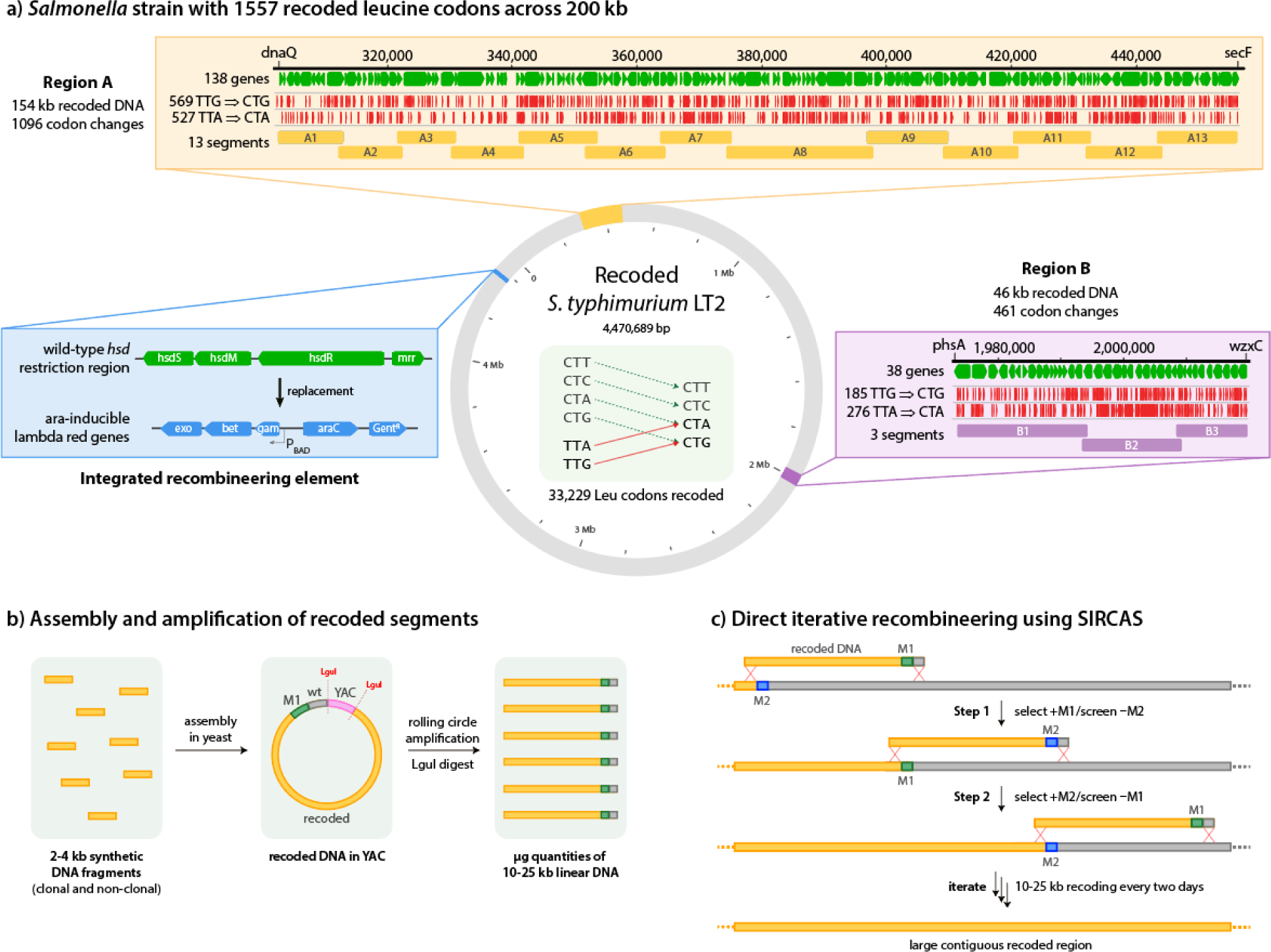
Rapid genome recoding by SIRCAS. a) From an *in silico* design of recoded *S. typhimurium* (grey), regions A and B (yellow/purple) were recoded in two strains, then combined by conjugative assembly. The integrated recombineering element (blue), inserted at the *hsd* restriction locus of *S. typhimurium*, is an arabinose-inducible lambda red cassette that enhances homologous replacement of wild-type genomic DNA with synthetic fragments. b) Recoded 10-25 kb segments were assembled from synthetic DNA fragments in yeast, amplified by rolling circle and linearized. Each segment contains a selection marker and homology flanks for integration. c) Accumulated recoding by SIRCAS involves iterative recombineering of recoded DNA segments. Each cycle takes two days to complete. Swapped selection markers provide a simple readout for successful recoding.

To demonstrate the power of our recoding approach, we replaced leucine codons due to their high frequency and redundancy in the *Salmonella* genome^14^. We computationally generated a recoded *S. typhimurium* genome in which all 33229 TTA/TTG leucine codons were replaced with synonymous CTA/CTG codons (Supplementary Note 1). In addition to recoding, 387 kb of pseudogenes, mobile elements and pathogenicity islands were removed to reduce genome size and instability, and 754 restriction sites for the enzyme LguI were removed to facilitate downstream cloning. From this design, we constructed 16 recoded segments (A1-A13 and B1-B3) constituting a 154 kb region A and a 46 kb region B (Figure 1a, Supplementary Note 2 and Data 1). Each segment contained 10-25 kb of recoded DNA, a selection marker, and 1 kb flanking homology regions for integration. The 10-25 kb size range was chosen to simplify construction, decrease the likelihood of unwanted internal recombination events, and minimize the cost of fixing an error in any one segment. Each segment was assembled in yeast from commercially synthesized 2-4 kb DNA fragments (Figure 1b)^15^. SIRCAS uses a marker swapping approach alternating between chloramphenicol and kanamycin selection (Figure 1c) for a simple phenotypic readout, similar to the strategy for building chromosomes in *S. cerevisiae^4^.*

We bypassed the use of bacterial plasmids and associated cloning steps by amplifying each recoded segment directly from yeast using rolling circle amplification^16^ and linearizing the resulting concatemer by LguI digestion to obtain microgram quantities of DNA for direct integration (Figure 1b). Additionally, carrying each segment on a bacterial plasmid would have required additional negative selection against the plasmid backbone to distinguish between the desired integration event vs the plasmid simply existing as an extrachromosomal replicative element^3,6^.

To create a *S. typhimurium* strain with high recombination efficiency, we constructed an integrated recombineering element (IRE) containing the lambda red genes under arabinose-inducible control^9^ (Supplementary Note 3). Targeted integration of the IRE to the *hsd* locus simultaneously removed the native *hsd* restriction system^17^ which could otherwise have impeded transformation (Figure 1a).

We successfully assembled 200 kb of recoded genome in a single strain of *S. typhimurium.* Hundreds of marker positive colonies were typically obtained after each round SIRCAS (Supplementary Figure 1). No colonies were obtained when arabinose or DNA was omitted. The proportion of colonies with correctly swapped markers ranged between 3-41% (median 14%, Supplementary Figure 1 and Table 1), presumably due to differences in marker integration locus^18^, as well as the size and content of the recoded DNA. Between each round of SIRCAS, we briefly checked for unwanted internal recombination events with Sanger sequencing. In 83% of all Sanger sequenced colonies over 16 rounds of SIRCAS, unwanted recombination events were not observed (Supplementary Table 2). Each round of SIRCAS required two days to complete (Supplementary Table 3), and only correct recombinants were used for further rounds of SIRCAS. After recoded regions A and B were assembled in two *S. typhimurium* strains, a conjugation step transferred region A into the strain carrying region B^19^ (Supplementary Note 4). Whole genome sequencing of the final strain confirmed perfect recombination of each recoded segment across the entire 200 kb.

We analyzed the error rates of SIRCAS, comparing commercial synthetic DNA obtained from clonal and non-clonal sources (Supplementary Note 5 and Data 1). Within the 156 kb of recoded regions that was written with clonal sequence-verified DNA, an overall error rate of 1 in 20000 was observed (7 point mutations and one leucine codon reversion in 200 kb). In comparison, Ostrov *et al.*^3^ reported an error rate of 1 in 5000 (an average of 9.7 mutations and 0.6 codon reversions in 50 kb). For the 44 kb of recoded regions written using non-clonal DNA, an overall error rate of 1 in 860 was found (51 errors, primarily single point deletions). Use of clonal DNA for SIRCAS is therefore preferable, producing an error rate that is competitive with that of other genome recoding methods.

Despite the vast number of changes introduced by recoding, doubling times were similar across all strains at 37 °C in LB (Figure 2, Supplementary Note 6). This result demonstrates that the *Salmonella* genome is amenable to large-scale recoding. Sequencing did not reveal any compensatory mutations in non-recoded regions of the genome (Supplementary Data 1).

**Figure 2.**
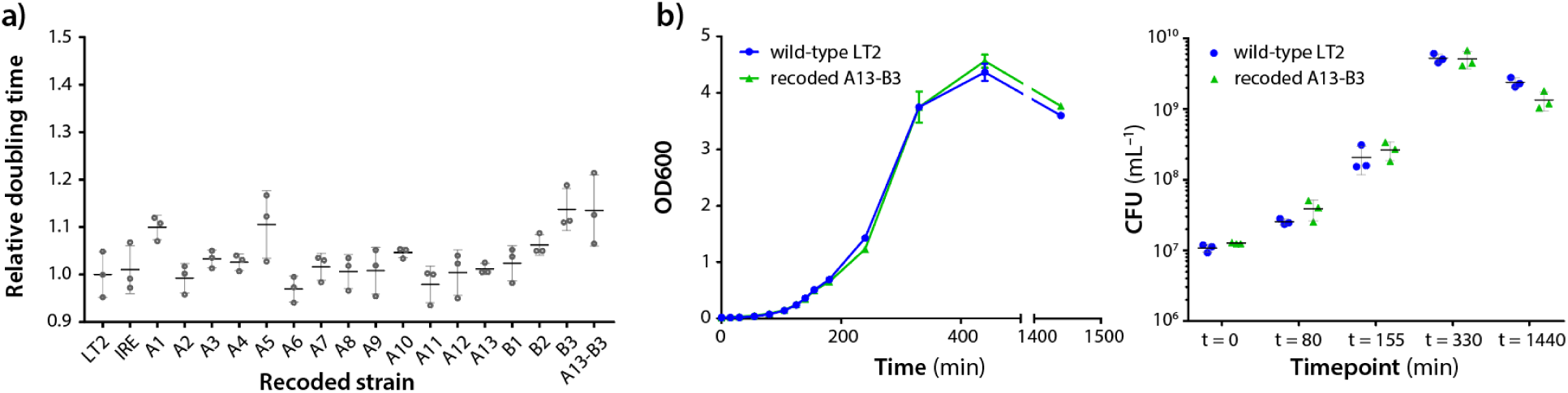
Growth rate of recoded strains. a) Doubling times at 37 °C in LB were similar for all the recoded strains over 16 rounds of SIRCAS. The strain nomenclature ‘A_’ refers to the strain containing all recoded segments from A1 up to A_. Each data point is the mean of three technical replicates, and the error bars represent the standard deviation of three biological replicates. b) Comparable growth curves and colony forming unit (CFU) measurements were observed for wild type *S. typhimurium* LT2 and the final recoded strain A13-B3. Each data point in the growth curve represents the average of three biological replicates, while each CFU data point is a single biological replicate. Errors bars represent then standard deviation.

In summary, SIRCAS is a rapid genome recoding method that does not have site-specific enzyme requirements. By integrating 20 kb recoded segments every two days, it is possible in one month to recode 300 kb in a single strain and an entire 4.5 Mb *Salmonella* genome by SIRCAS in 16 parallel strains. With this recoding method, we can achieve genetic containment^20^ of engineered *Salmonella* for therapeutic applications, and create strains for expressing proteins containing unnatural amino acids^21^. Beyond recoding, the use of SIRCAS to rapidly and precisely rewrite entire gene clusters enables interrogation of *Salmonella* genetics on a scale that is not possible with traditional techniques. Importantly, SIRCAS is not limited to *Salmonella,* but should be applicable in any recombineering compatible host, and has utility for many large-scale genome engineering applications, such as rapid integration of entire *de novo* designed biosynthetic pathways into industrial production strains.

## METHODS

Methods are described in the supplementary methods file.

## AUTHOR CONTRIBUTIONS

The overall study was conceived by PAS and JCW. Experimental design was conducted by YHL, FS, JK, MAPK and AR. The computational design of the recoded genome and other computational analyses were performed by MAPK, AR and YHL. Assembly and RCA of DNA constructs was performed by YHL, FS, CAH, ES and DL. The protocols for DNA assembly and RCA were established through guidance and troubleshooting from MTW, DGG and YAC. Genomic integration (SIRCAS) was conducted by YHL and CAH. Conjugation experiments were performed by JK. Sequence analysis was conducted by YHL and FS. Growth data for recoded strains was obtained by YHL. All authors contributed to writing the manuscript.

## ACKNOWLEDGEMENTS

We thank Eric Kofoid and Natalie Duleba from the laboratory of Prof. John Roth (Department of Microbiology, University of California Davis) for the kind gift of the recombineering *Salmonella* strain TT22971 containing plasmid pKD46. We also thank Prof. George Church (Department of Genetics, Harvard Medical School) and members of his laboratory for general discussions about genome recoding.

## REFERENCES

1. Boeke, J. D. et al. Science 353, 126–127 (2016).

2. Lajoie, M. J. et al. Science 342, 357–360 (2013).

3. Ostrov, N. et al. Science 353, 819–822 (2016).

4. Annaluru, N. et al. Science 344, 55–58 (2014).

5. Hutchison, C. A. et al. Science 351, aad6253 (2016).

6. Wang, K. et al. Nature 539, 59–64 (2016).

7. Wang, H. H. et al. Nature 460, 894–898 (2009).

8. Nyerges, Á. et al. Proc. Natl. Acad. Sci. U.S.A. 113, 2502–2507 (2016).

9. Datsenko, K. A. & Wanner, B. L. Proc. Natl. Acad. Sci. U.S.A. 97, 6640–6645 (2000).

10. Yu, D. et al. Proc. Natl. Acad. Sci. U.S.A. 97, 5978–5983 (2000).

11. Kotula, J. W. et al. Proc. Natl. Acad. Sci. U.S.A. 111, 4838–4843 (2014).

12. Zheng, J. H. et al. Sci. Transl. Med. 9, eaak9537 (2017).

13. Kong, W., Brovold, M., Koeneman, B. A., Clark-Curtiss, J. & Curtiss, R. Proc. Natl. Acad. Sci. U.S.A. 109, 19414–19419 (2012).

14. McClelland, M. et al. Nature. 413, 852–856 (2001).

15. Gibson, D. G. et al. Proc. Natl. Acad. Sci. U.S.A. 105, 20404–20409 (2008).

16. Dean, F. B., Nelson, J. R., Giesler, T. L. &Lasken, R. S. Genome Res. 11, 1095–1099 (2001).

17. Colson, C. & Van Pel, A. Mol. Gen. Genet. 129, 325–337 (1974).

18. Englaender, J. A. et al. ACS Synth. Biol. Article ASAP (2017). doi:10.1021/acssynbio.6b00350

19. Isaacs, F. J. et al. Science 333, 348–353 (2011).

20. Ma, N. J. & Isaacs, F. J. Cell Syst. 3, 199–207 (2016).

21. Kipper, K. et al. ACS Synth. Biol. 6, 233–255 (2017).

